# Conservation genomics of desert tortoises (*Gopherus agassizii*) in the Colorado Desert

**DOI:** 10.64898/2025.12.08.692717

**Authors:** W. Bryan Jennings, Jeffrey E. Lovich, Kristy L. Cummings, Shellie R. Puffer, Joshua R. Ennen, Mickey Agha, Kathleen D. Brundige, Christopher R. Tracy

## Abstract

Management units (MUs) are important to conserving species of conservation concern. Although the MU definition is simple (i.e., a demographically independent population), identifying MUs in practice is difficult because investigators must choose a recent (i.e., last few generations) migration rate threshold. One suggested MU criterion is *m* < 0.10, where *m* is the proportion of a subpopulation comprised of recent migrants. However, a more recent study found that *m* as low as 0.02 (i.e., one migrant per generation) can cause estimates of linkage disequilibrium effective population size (*N*_eLD_) to underestimate a subpopulation’s true *N*_e_. Here, we used genome-wide SNP data obtained from 40 tortoises and an MU criterion of *m* < 0.02 to define MUs within the Colorado Desert population of the Mojave desert tortoise (*Gopherus agassizii*)—an endangered species in California and a species in the USA protected under the Endangered Species Act. Based on our results we defined the Mesa and Deep Canyon subpopulations as MUs even though the latter received at least two recent immigrants. Migrant(s) from the Shavers Valley subpopulation were likely translocated by humans, whereas migrant(s) from the Mesa subpopulation could have been human-mediated or natural migrants. Initial analyses suggest that the Shavers Valley subpopulation may be an MU, but our low sample size renders this result inconclusive. The local *N*_e_ estimates for the Mesa and Deep Canyon subpopulations—5 and 13 adults, respectively—are below the minimum sizes according to the 50/500 rule and thus they may be candidates for genetic rescue.

## Introduction

Populations of the Mojave desert tortoise (*Gopherus agassizii*) have been declining for many decades due to diseases and myriad anthropogenic activities such as habitat disturbances, livestock grazing, mining, urban and agricultural development, roads and utility corridors, recreational vehicle use, solar energy development, and military training activities (U.S. Fish and Wildlife Service [USFWS] 1994; Lovich and Bainbridge 1999; Edwards et al. 2004; Lovich and Ennen 2011; Berry and Murphy 2019). In 1990, the USFWS listed this species as a threatened species under the Endangered Species Act (USFWS 1990, 1994). Despite efforts to recover populations of the Mojave desert tortoise, this species continues to decline (USFWS 2011; Allison and McLuckie 2018; Berry and Murphy 2019; Berry et al. 2021; Zylstra et al. 2023). The decline has become so pronounced, that the California Department of Wildlife changed the designation of this species in the state from a threatened to an endangered species in 2024.

Studies aimed at elucidating the population genetic structure of the Mojave desert tortoise represent an important part of on-going efforts to save this wide-ranging species (Lamb et al. 1989; Murphy et al. 2007; Sánchez-Ramirez et al. 2018; Haggerty and Tracy 2010; Lovich et al. 2020). Critical tasks ahead include the delineation of conservation units (CUs), determining levels of connectivity among them, and estimating the genetic diversity of each CU (Waples and Gaggiotti 2006; Palsbøll et al. 2007; Palstra and Ruzzante 2008; Funk et al. 2012; Hohenlohe et al. 2021; Waples 2024).

The authors of the *Desert Tortoise Recovery Plan* (USFWS 1994) used available genetic, ecological, behavioral, and morphological evidence to define six “recovery units,” which according to Murphy et al. (2007) equated to CUs at the level of “Evolutionary Significant Units” (or ESUs) and “Distinct Population Segments” (or DPS; Ryder 1986; U.S. Department of the Interior and U.S. Department of Commerce; Waples 1991, 1998). However, because the genetic evidence (i.e., mitochondrial DNA restriction fragment length polymorphisms data; Lamb et al. 1989) underpinning these recovery unit designations had low resolving power, Murphy et al. (2007) advocated the use of more powerful population genetic data types available at the time (i.e., microsatellites) to corroborate the recovery units as ESUs/DPSs. Subsequent studies of the Mojave desert tortoise based on mtDNA sequences and microsatellites (Murphy et al. 2007; Lovich et al. 2020), microsatellites only (Haggerty and Tracy 2010), genome-wide Single Nucleotide Polymorphisms or “SNPs” (Sánchez-Ramirez et al. 2018) largely supported the recognition of the original USFWS recovery units as being ESUs and DPSs. Despite these advances, we still lack sufficient knowledge about the fine-scale population genetic structure within these recovery units/DPSs/ESUs.

Progress has been made towards understanding the structure within one of the desert tortoise recovery units, the Eastern Colorado Desert or “EC” recovery unit in southeastern California. This recovery unit was thought to only comprise tortoise subpopulation(s) east of the Coachella Valley, distributed among the Chocolate Mountains, Cottonwood Mountains, Eagle Mountains, Orocopia Mountains, Chuckwalla Mountains, and intervening valleys (USFWS 1994; Murphy et al. 2007; Haggerty and Tracy 2010; Sánchez-Ramirez et al. 2018). However, a later study by Lovich et al. (2020) based on mtDNA sequences and microsatellites showed that an apparently isolated subpopulation in the foothills of the San Bernardino Mountains in the western part of the Coachella Valley near Whitewater, which they referred to as the “Mesa subpopulation,” was most closely related to tortoises found just east of the Coachella Valley between the Cottonwood and Orocopia Mountains in Shavers Valley, hereafter the “Shavers Valley subpopulation.” The closest known desert tortoise aggregation to Mesa is about 10 km to the northeast in the foothills of the Little San Bernardino Mountains (J. Lovich pers. obser). However, as those tortoises occurred within the confines of the Coachella Valley, they were likely part of the Mesa subpopulation before human developments in the area apparently isolated them. Unfortunately, the Lovich et al. (2020) study did not include samples from another unstudied Colorado Desert subpopulation located on the desert slopes of the Santa Rosa Mountains near Palm Desert, hereafter “Deep Canyon subpopulation” (Lovich et al. 2015, 2018, 2020; Berry and Murphy 2019). Given the vast distances between these three EC subpopulations as well as many intervening barriers to dispersal including lack of tortoise habitat and human infrastructure such as Interstate 10, urban development, and concentrated agricultural developments (Figure 1), it is likely that these subpopulations have been isolated from each other for at least 50 years, if not over longer evolutionary time.

**Figure 1.**
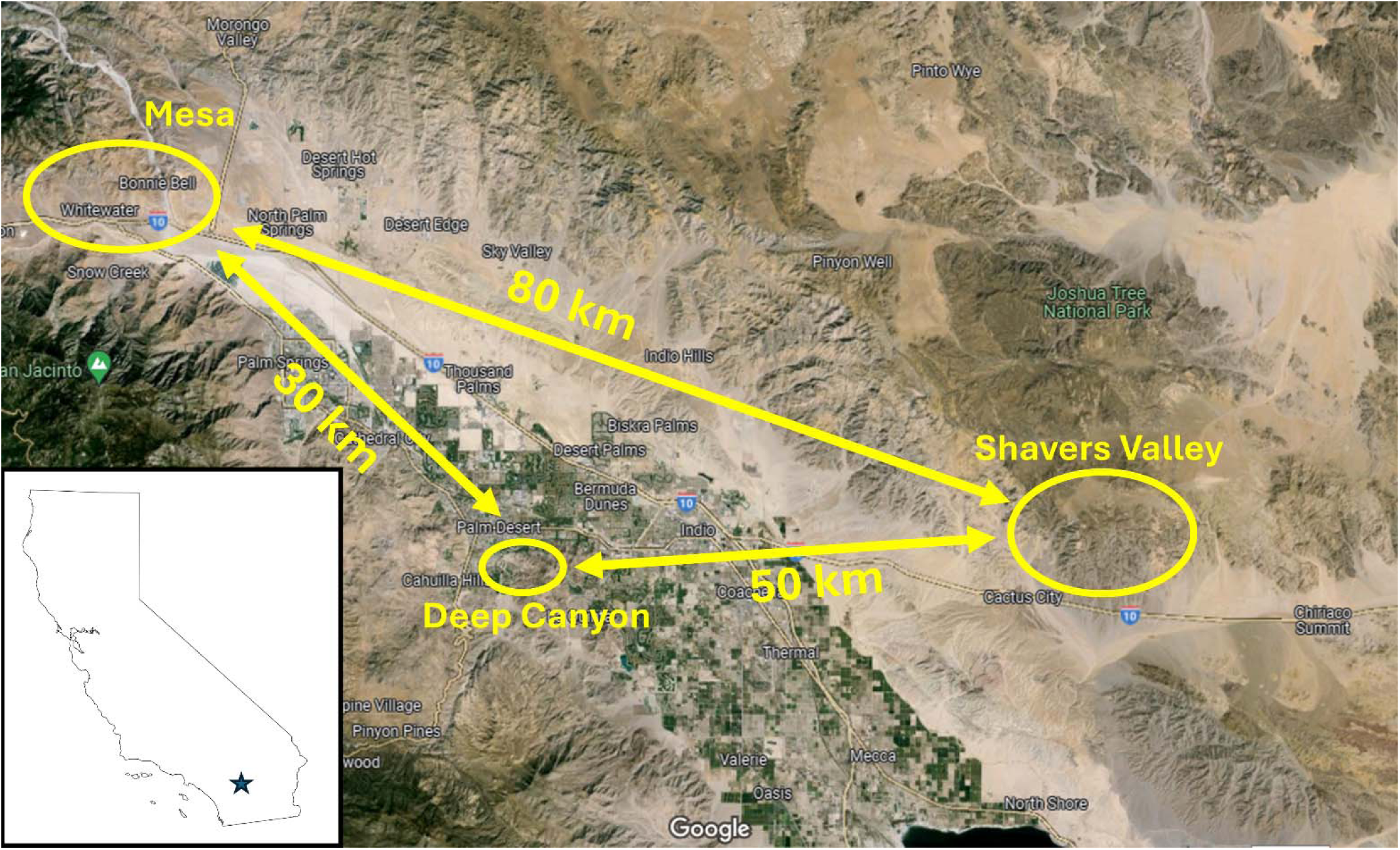
Map showing the locations of the Deep Canyon (33.65 lat, -116.38 long), Mesa (33.93, -116.62), and Shavers Valley (33.67, -115.79) study sites. The approximate boundaries of each site are delineated by yellow ellipses and straight-line distances between sites are shown by the yellow lines. Image source: Google Earth. Black star in California state map shows the location of the study region.

A population genomics approach that can reveal levels of recent genetic divergence between subpopulations is needed to determine if these tortoise subpopulations are demographically independent of each other and thus represent management units or “MUs” (Palsbøll et al. 2007; Funk et al. 2012; Hohenlohe et al. 2021). MUs are the CUs of main interest to land and wildlife managers because these population units are central to the short- and long-term management and conservation of natural populations (Palsbøll et al. 2007; Funk et al. 2012; Hohenlohe et al. 2021). One approach to delineating MUs is to estimate the recent (i.e., over the past few generations) migration rates between two subpopulations; if the immigration rate into a subpopulation is less than an *a priori* threshold value, then that subpopulation can be considered as an MU (Waples and Gaggiotti 2006; Palsbøll et al. 2007). The parameter *m* is defined as the fraction of a subpopulation comprised of immigrants from other subpopulations (Wilson and Rannala 2003). As Waples and Gaggiotti (2006) pointed out, the chief difficulty with this approach lies in the choice of threshold for *m*. Hastings (1993) suggested that an *m* < 0.10 could be used as a criterion for demographically independent subpopulations, whereas Palsbøll et al. (2007) suggested that thresholds might be best determined using individual-based population computer models. An alternative approach to delineating MUs was suggested by Funk et al. (2012), who advocated the use of model-based clustering analyses of presumably neutral SNPs to delineate MUs. Applications of this latter approach include the studies of Barbosa et al. (2018) and Forester et al. (2022a), though the latter study also used several genetics-based methods in conjunction with environmental data and information on dispersal capacity to delineate MUs in their target species. Given the importance of MUs to conservation efforts and the lack of agreement on how to delineate these units, it is imperative that more work be done to establish guidelines for the future.

Another important aspect of MUs is that once they have been delineated, the local contemporary effective population size (*N_e_*) of each MU can be estimated. *N_e_* is a key parameter in population genetics because it determines the rate of genetic drift across the genome (Waples 2022). It can be defined as a quantity based on three demographic aspects of real populations: 1) the number of potential parents; 2) the mean number of offspring per parent; and 3) the variance in the number of offspring per parent (Waples 2022). This parameter is also critically important in conservation studies because it is tied to the genetic health and evolutionary potential of species (Franklin 1980; Mace and Lande 1991; Frankham 1995; Palstra and Ruzzante 2008; Funk et al. 2019; Hoban et al. 2020, 2021, 2024; Forester et al. 2022b).

The most common method for estimating *N*_e_ using genome-wide SNP data is the single-sample linkage disequilibrium (LD) approach, hereafter referred to as the “*N*_eLD_ method,” as opposed to the older two-sample “temporal method” (Waples and Do 2009; Do et al. 2014; Waples 2023; Santiago et al. 2023). *N*_eLD_ is used as a proxy for the target *N*_e_ parameters in conservation studies, which are the inbreeding effective size (*N*_eI_) and additive variance effective size (*N*_eAV_; Waples and Do 2009, Ryman et al. 2019; Waples 2023). This is important because the venerable “50/500 rule” in conservation genetics dictates that populations having an *N*_eI_ < 50 will have unacceptably high inbreeding rates while populations with *N*_eAV_ < 500 will lose additive genetic diversity and hence become less able to adapt to changing environmental conditions over the long term (Franklin 1980; Jamieson and Allendorf 2012; Ryman et al. 2019; Hoban et al. 2021; but see Frankham et al. 2014). When subpopulations are demographically independent of other subpopulations and are of constant size over a long period of time, *N*_eLD_ will be equivalent to *N*_eI_ and *N*_eAV_ (Ryman et al. 2019). Although simulation studies suggested that estimates of *N*_eLD_ are robust to migration levels up to around *m* = 0.10 (Waples 2010; Waples and England 2011), migration rates as low as one migrant per generation (*m* = 0.02) can, over time, still cause the local *N*_e_ for a subpopulation to converge upon the larger metapopulation *N*_e_—meaning that local *N*_eLD_ estimates will increasingly underestimate *N*_eI_ and *N*_eAV_ over time (Ryman et al. 2019). It is therefore essential that researchers carefully interpret their local *N*_eLD_ estimates considering what is known about gene flow levels between subpopulations (Waples 2010, 2022, 2023; Waples and England 2011; Ryman et al. 2019).

Here, we evaluated the population genetic structure and conservation status of Mojave desert tortoise subpopulations within the Colorado Desert region of California (a western extension of the Sonoran Desert named for its proximity to the Colorado River) using thousands of genome-wide SNPs and recently developed bioinformatics tools. Our goals were to: 1) determine if the Mesa, Deep Canyon, and Shavers Valley tortoise subpopulations represent three MUs or form a larger metapopulation; 2) estimate *N*_eLD_ for each MU or metapopulation; 3) evaluate the robustness of our *N*_eLD_ estimates by integrating data for the census population sizes of adults (*N*) obtained from mark-recapture studies; and 4) determine the short-term risk of inbreeding and long-term vulnerability to reduced evolutionary potential for each closed population unit. Our findings could have important implications for the management and conservation of subpopulations of this species.

## Materials and methods

### Genetic Samples

Fresh blood samples were obtained from 40 free-ranging adult tortoises. Nine of the samples were taken from the Deep Canyon subpopulation, 23 samples were obtained from the Mesa subpopulation, and eight samples were taken from the Shavers Valley subpopulation.

Approximately 500 μL of fresh blood was taken from each tortoise and placed in a sterile 2 mL cryopreservation tube containing ∼500 μL of 100% ethanol before being kept cold on ice or in a - 80°C freezer until used in this study. All laboratory work was conducted in the Genomics Core Facility at the University of California, Riverside. Purified genomic DNA was extracted from each blood sample using a Qiagen DNA Easy Blood and Tissue Kit (Hilden, Germany) with an RNase A treatment following the manufacturer’s instructions. The purified DNA samples were then shipped to Floragenex, Inc. (Beaverton, Oregon) for double-digest restriction site (ddRADSeq) library preparation (Peterson et al. 2012) and sequencing. Libraries were made using PstI and MseI enzymes with the added step of using a +2 primer (with the sequence +CC) that extends 2 base pairs (bp) past the MseI cut site into the gDNA to further reduce the number of loci. Libraries were sequenced on an Illumina Novaseq 6000 platform in an S1 flow cell to generate 100 bp single end reads. All data analyses except some (indicated below) were run on the high-performance computing cluster at the University of California, Riverside (UCR HPCC).

### Bioinformatics processing of raw data

We assembled the raw fastq sequence reads into *de novo* ddRADSeq loci using the *iPyrad* Python pipeline v0.9.84 (Eaton and Overcast 2020). Initially, this program demultiplexes the samples according to their unique barcode sequences while using a strict filter for adapter and primer sequences. The following parameters were used in the parameter file for the assembly: clustering threshold of 0.91, minimum depth of ten for base calling and majority rule base calling, sites with Phred quality scores below 33% (99.95%) were converted to ‘N’ characters, reads with >5 N’s were filtered out, a maximum of 50% heterozygous sites per locus were allowed, a maximum of 5% uncalled bases in consensus sequences were allowed, a maximum of 5% heterozygotes in consensus sequencies were allowed, and we set the minimum number of samples output to 38 (95%). Filtered data were converted into various file formats depending upon the population genetics program used (see below). All raw genomic sequences and datafiles used in analyses are found in the Dryad repository under DOI: xxxxxxxxx.

### Population genetic structure

#### Analyses of genetic structure using model-based approaches

We used the software package *fastStructure* v.1.0 9 (Raj et al. 2014) to investigate population genetic structure of our three study subpopulations. We chose this model-based clustering program because it is more computationally efficient than its predecessor *STRUCTURE* (Pritchard et al. 2000) and because it can detect weak population structure better than other comparable programs (Raj et al. 2014). *fastStructure* implements a variational Bayesian framework with two options for priors. The “simple prior” is computationally fast but may not resolve subtle population structure, whereas the “logistic prior” has a larger computational burden but tends to resolve subtle structure better than the simple prior. The program generates two heuristic scores to choose model complexity or *K*, the number of distinct population clusters: the first heuristic, *K*\*, is chosen to maximize the marginal likelihood of the data, while the other, *K*_Ø_c, equates the minimum number of populations as the number of relevant model parameters that have a cumulative ancestry contribution of each least 99.99% (Raj et al. 2014). When there is strong structure and a simple prior is used, both *K*\* and *K* c are expected to match *K*. However, when there is very weak structure in the data, the former metric severely underestimates *K* while the latter slightly overestimates *K* (Raj et al. 2014). Thus, in cases when a simple prior yields different values for *K*\* and *K* c, the authors recommended using a logistic prior. One problem with the logistic prior is that *K*\* can overestimate *K* due to model overfitting (Raj et al. 2014). In such cases *K*_Ø_c may better reflect *K*. *fastStructure* requires the data input file to be in either str (*STRUCTURE*) or bed formats. We constructed a data file in str format using the following steps. First, we filtered the vcf file output from *iPyrad* using *VCFTools* v0.1.13 (Danecek et al. 2011) so that the SNP data met the population genetic assumptions of *fastStructure*. To ensure that each SNP was unlinked from other sampled SNPs (Waples et al. 2022) we set the thinning parameter to 107 bp to limit one SNP per locus. We also filtered the data to keep only biallelic SNPs using the following settings: - -min-alleles 2 and - -max-alleles 2. Given the potentially adverse impacts of singletons (caused by natural base substitutions and sequencing errors) on *STRUCTURE*-type analyses, it is advisable to exclude singletons from analyses (Linck and Battey 2019). We therefore filtered out singletons by setting the minor allele frequency (MAF) to 0.05. The --max-missing parameter was set to 0.95. The filtered vcf file was then converted into a str file by using the vcfR library (Knaus and Grünwald 2017) in the *R* statistical package (version 4.3.0, R Development Core Team). All *R* analyses were performed on a Windows 10 machine. We ran *fastStructure* with *K* = 1 to *K* = 5 for both simple and logistic priors. Lastly, we used the *chooseK.py* script to output estimates for *K*\* and *K* c.

#### Analyses of genetic structure using non-model-based approaches

Given that model-based programs like *STRUCTURE* contain assumptions such as the type of population subdivision, best practices dictate that non-model-based clustering analyses such as PCA (Principal Components Analysis) and DAPC (Discriminant Analysis of Principal Components) should also be conducted to corroborate results obtained from model-based clustering analyses (Jombart et al. 2010; Linck and Battey 2019). We therefore analyzed the same filtered vcf file we generated for the *fastStructure* analyses using PCA and DAPC as implemented in the *R* package *Adegenet* 2.0-0 (Jombart et al. 2008, 2010). After running the PCA analysis, we visualized the number of genetically distinctive clusters by plotting the first two principal components (PCs). However, two disadvantages of PCA compared to DAPC is that it typically only considers a small percentage of the variation in the data (i.e., from PC axes 1 and 2) and PCA by itself cannot be used to objectively determine *K* (Jombart et al. 2010). In contrast, DAPC considers most of the variance in the data and it can cluster individuals into genetically distinctive groups without any previous knowledge of where they were sampled. The DAPC analysis comprised several steps. First, we used PCA to transform the data into uncorrelated variables. Next, a cross-validation procedure was used to determine the optimal number of PCs to retain for the discriminant analysis (DA) step. The *find.clusters* function was then used for *K* = 1 to *K* = 6 to find the optimal *K* value for the data.

#### Estimation of recent migration rates

We used the software package *BA3-SNPs* (Mussmann et al. 2019) to estimate recent migration rates (*m*) between the three tortoise subpopulations. *BA3-SNPs* is a package that adapted the Bayesian Markov Chain Monte Carlo (MCMC) program *BayesAss* (Wilson and Rannala 2003) for use with genome-wide SNP datasets. Only the previous two generations of migrant ancestry are used to estimate *m* because older generations are statistically indistinguishable from non-migrants (Wilson and Rannala 2003). *VCFTools* was used to filter the original vcf file so that the new file contained biallelic SNPs, one SNP/locus, and had no missing data to satisfy the assumptions of the analysis and requirements of the program. We then converted the filtered vcf file into the required immanc file format using conversion scripts included in the *BA3-SNPs* package. The autotune script conducted a short preliminary run with the objective of identifying suitable starting values for the three mixing parameters. The *BA3-SNPs* program was run with the following settings specified in the command line: 20 million generations with the first 15 million generations discarded as burn-in, sample interval of 2,000 generations, and the mixing parameters M, A, and F were set to 0.2125, 0.5500, and 0.0250, respectively. Three independent runs of *BA3-SNPs*, each using a different random seed, were conducted to ensure that parameter estimates achieved convergence. Each *BA3-SNPs* run output six migration estimates, which included two (bi-directional) migration rates for each pair of subpopulations, along with the standard deviation of the marginal posterior distribution for each estimate. The program also infers the individual migrant histories; that is, each individual was classified as a non-migrant, a first-generation migrant, or a second-generation migrant. We also computed Bayesian 95% credible intervals for all estimates of *m* as the posterior means ± 1.96 × standard deviations. We considered a subpopulation to be an MU (i.e., a demographically independent subpopulation) if the posterior mean *m* was less than 0.02 because this level of migration is not expected to impact estimation of local *N_e_*.

#### Estimation of LD-based contemporary effective population sizes

We first attempted to analyze our data using the program *NeLD* (Waples and Do 2009; Do et al. 2014) but we could not obtain results possibly due to our low sample sizes for two of the subpopulations. We therefore used the program *CurrentNe* (Santiago et al. 2023) to obtain *N*_eLD_ estimates for the Deep Canyon, Mesa, and Shavers Valley subpopulations. This program estimates *N*_eLD_ as the average effective population size over the past couple generations (i.e., the sampling generation and one generation before that). *CurrentNe* uses an artificial neural network to estimate contemporary *N*_eLD_ and associated 90% confidence intervals. We used the program *VCFTools* (Danecek et al. 2011) to prepare subpopulation-specific vcf files that were suitable for *CurrentNe*. These files were filtered the same way as before except for the following difference. *CurrentN*e assumes that the SNP allele frequencies in the data are representative of those found in the population and therefore rare alleles should not excluded from the vcf files using the MAF filter. The authors of this program determined that the main source of bias in *N*_eLD_ estimates is due to the type of mating system because this influences the number of full sibs in a sample.

Accordingly, the user must use *CurrentNe* to estimate the parameter k, and then re-run the program using the estimated k value to estimate *N_e_*. The k parameter, which generally ranges between 0 and 2, is the expected number of full siblings that a randomly selected individual will have among the reproducers in the population. However, we were unable to estimate k from the data in this manner and therefore a value for k had to be estimated using information about the desert tortoise mating system. The maximum value of k = 2 for a monogamous mating systems is not appropriate for desert tortoises because they have polygamous and polygynous mating systems (Davy et al. 2011; Mulder et al. 2017). Indeed, approximately 50% of clutches of tortoise eggs showed evidence of multiple paternity in desert tortoises (Pearse and Avise 2001).

Thus, we used k = 1 following the recommendations in the user manual for the program. The user must also input on the command line the number of chromosomes in the genome of the organism under study to run the program. We selected a value of 52 because the genome of desert tortoises contains 52 chromosomes (Stock 1972; Dowler and Bickham 1982).

## Results

### Sequencing data

The *iPyrad* pipeline processed a total of 370,401,711 raw sequence reads during the loci assembly process. There was an average of 9,260,043 reads per individual (range 4,650,712 to 17,060,360), which were assembled into an average of 11,068 loci per individual (range 6,340-11,453) across all 40 individuals.

### Population structure

#### Analyses of genetic structure using model-based approaches

Results from the *fastStructure* analysis based on a simple prior identified one to three distinct population clusters (*K*\* = 1 and *K* c = 3). The lack of agreement between the two heuristics based on the simple prior is evidence of weak population genetic structure. Because *K*\* and *K* c in this analysis are likely to be under- and overestimates of *K*, respectively, results based on the simple prior indirectly suggest there are *K* = 2 populations. Further evidence of weak population structure was provided by the *fastStructure* analyses based on a logistic prior, as the results suggested that there were between three and five genetically distinctive populations (*K*\* = 5 and *K*_Ø_c = 3). Given the apparent weak structure, the *K*\* = 5 result is likely to be an overestimate caused by model overfitting. Accordingly, results based on the logistic prior suggest that the optimal number of clusters is three. Visual inspection of ancestry proportions for *K* = 2 under both the simple and logistic models suggests that the tortoises from Deep Canyon are more similar genetically to the Shavers tortoises than either is to the Mesa subpopulation (Figure 2A, 2B). Under *K* = 3, both the simple and logistic models also suggest that the Deep Canyon and Shavers tortoises are very closely related to each other, while the Mesa tortoises appear to comprise two genetically distinct subgroups (Figure 2C, 2D).

**Figure 2.**
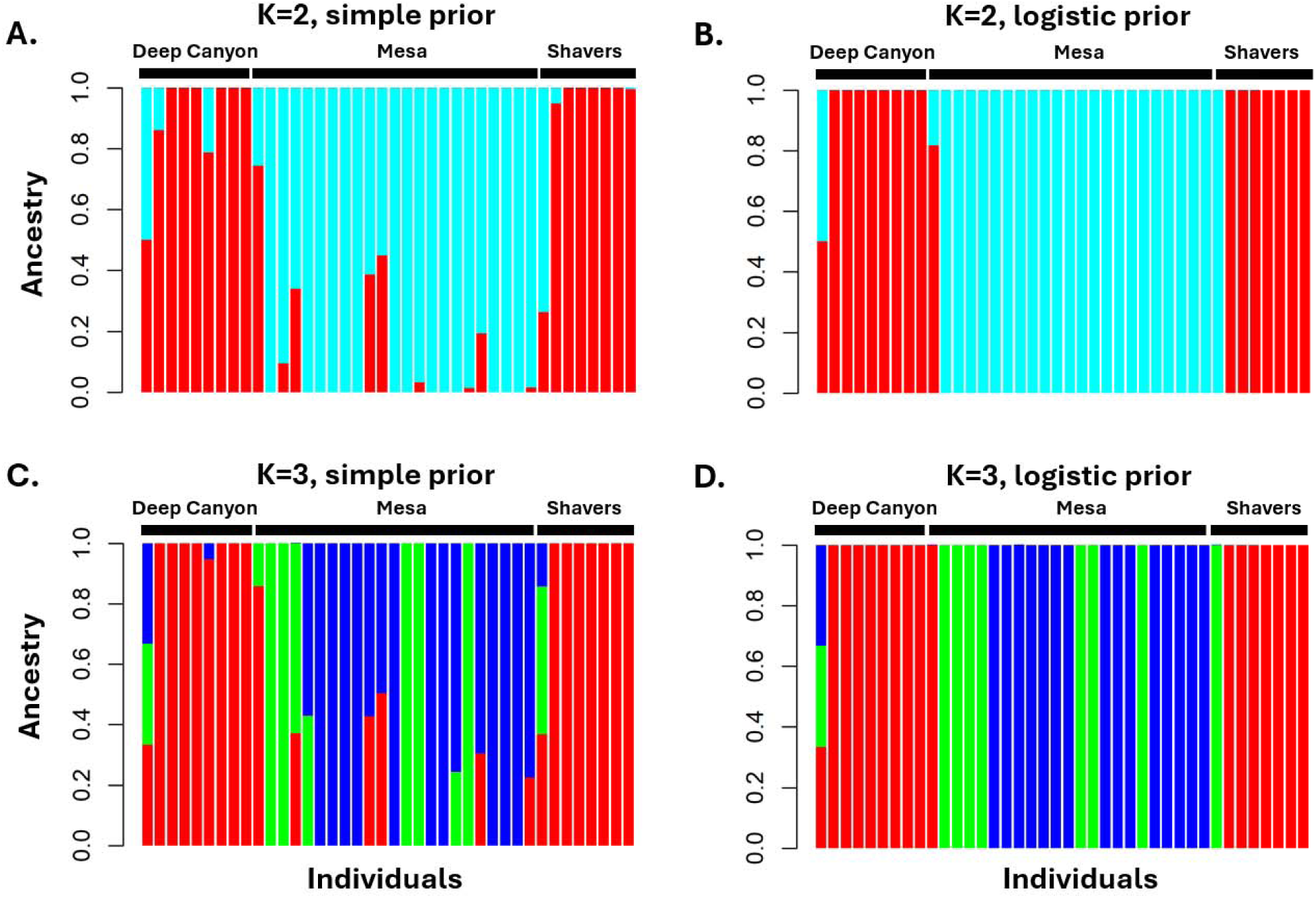
Admixture plots for tortoises in the Colorado Desert of California generated by the program *fastStructure*. Each vertical bar shows the color-coded ancestry proportions for an individual tortoise (n = 40 tortoises). (A) Admixture plot for *K* = 2 using a simple prior. (B) Admixture plot for *K* = 2 using a logistic prior. (C) Admixture plot for *K* = 3 using a simple prior. (D) Admixture plot for *K* = 3 using a logistic prior.

#### Analyses of genetic structure using non-model-based approaches

Results of the PCA analysis revealed two distinct clusters for the Mesa subpopulation (two individuals not clustering with the rest) and one cluster representing the Deep Canyon and Shavers Valley subpopulations (Figure 3). The plot also suggests that the Deep Canyon and Shavers Valley subpopulations are more genetically like each other than either one is to the Mesa tortoises, and it shows mostly non-overlapping points for the two subpopulations (Figure 3). Also evident in Figure 3 is at least two possible cases of admixture, as two Mesa tortoises clustered with the Deep Canyon tortoises, suggesting that the two individuals in question descended from ancestors of both subpopulations.

**Figure 3.**
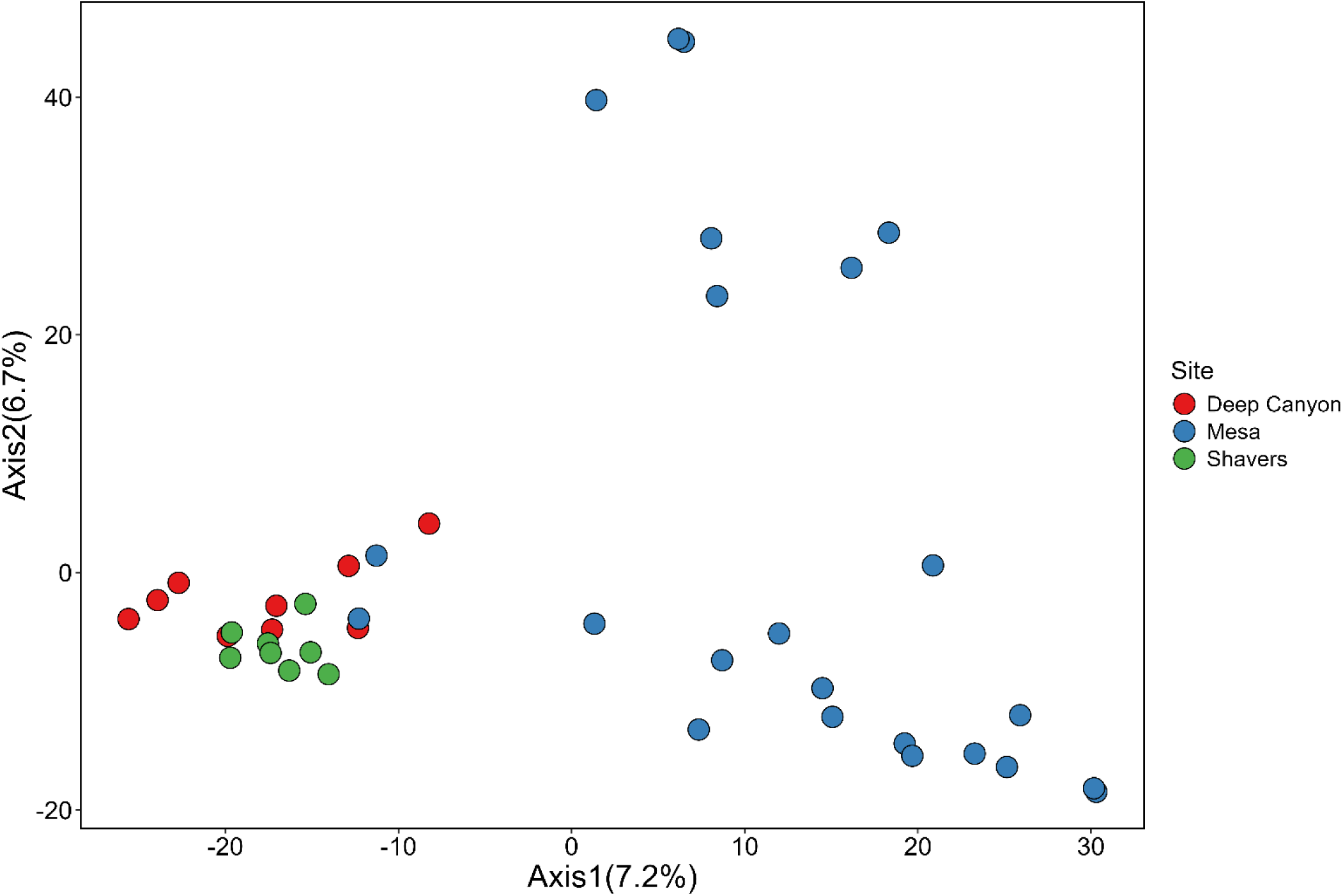
Scatterplot of principal component axes 1 and 2 from the Principal Components Analysis (PCA). Each point represents a sampled tortoise, whereas colors correspond to the Deep Canyon, Mesa, and Shavers Valley subpopulations.

The cross-validation procedure revealed that the first 20 principal components (PCs), which accounted for 68% of the variance, exhibited the lowest mean squared error. We retained these PCs for the DAPC analysis. The plot showing the Bayesian Information Criterion (BIC) value versus cluster number revealed the optimal number of clusters was three (Figure 4). The results of the DAPC showed that two discriminant functions were retained. When the first two PCs of the DAPC were plotted (Figure 5), the data showed the same three genetic clusters observed in the PCA plot, except that the points for the Deep Canyon and Shavers Valley subpopulations overlapped each other to a greater extent (compare Figures 3 and 5). Like the PCA plot, the Mesa subpopulation exhibits a relatively high amount of genetic divergence between two subgroups of tortoises and the DAPC plot shows two Mesa individuals that clustered with the Deep Canyon and Shavers tortoises (Figure 5).

**Figure 4.**
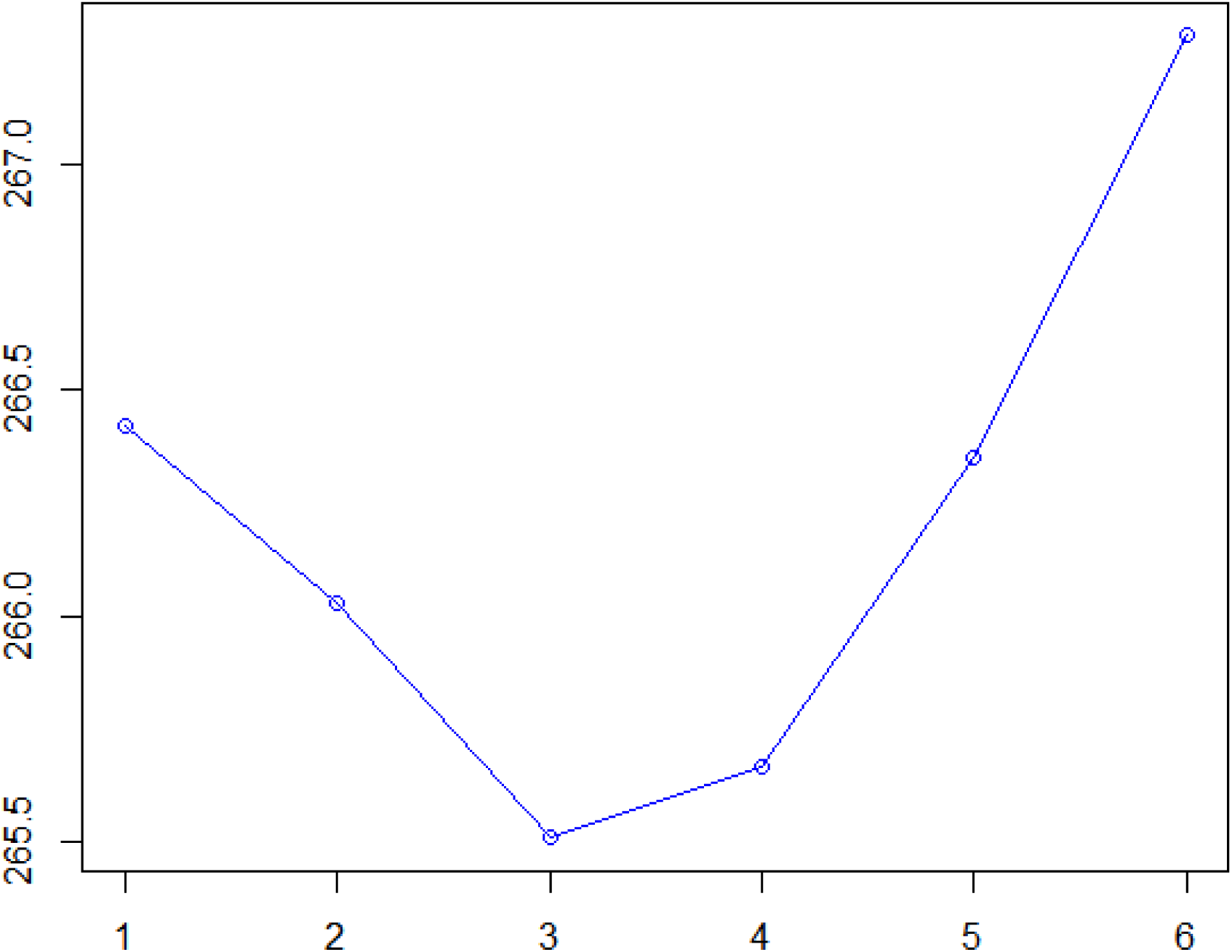
Bayesian Information Criterion (BIC) versus number of clusters in the Discriminant Analysis of Principal Components (DAPC) analysis. The BIC with the lowest value (i.e., “elbow” in the graph) represents the optimal number of clusters, *K*.

**Figure 5.**
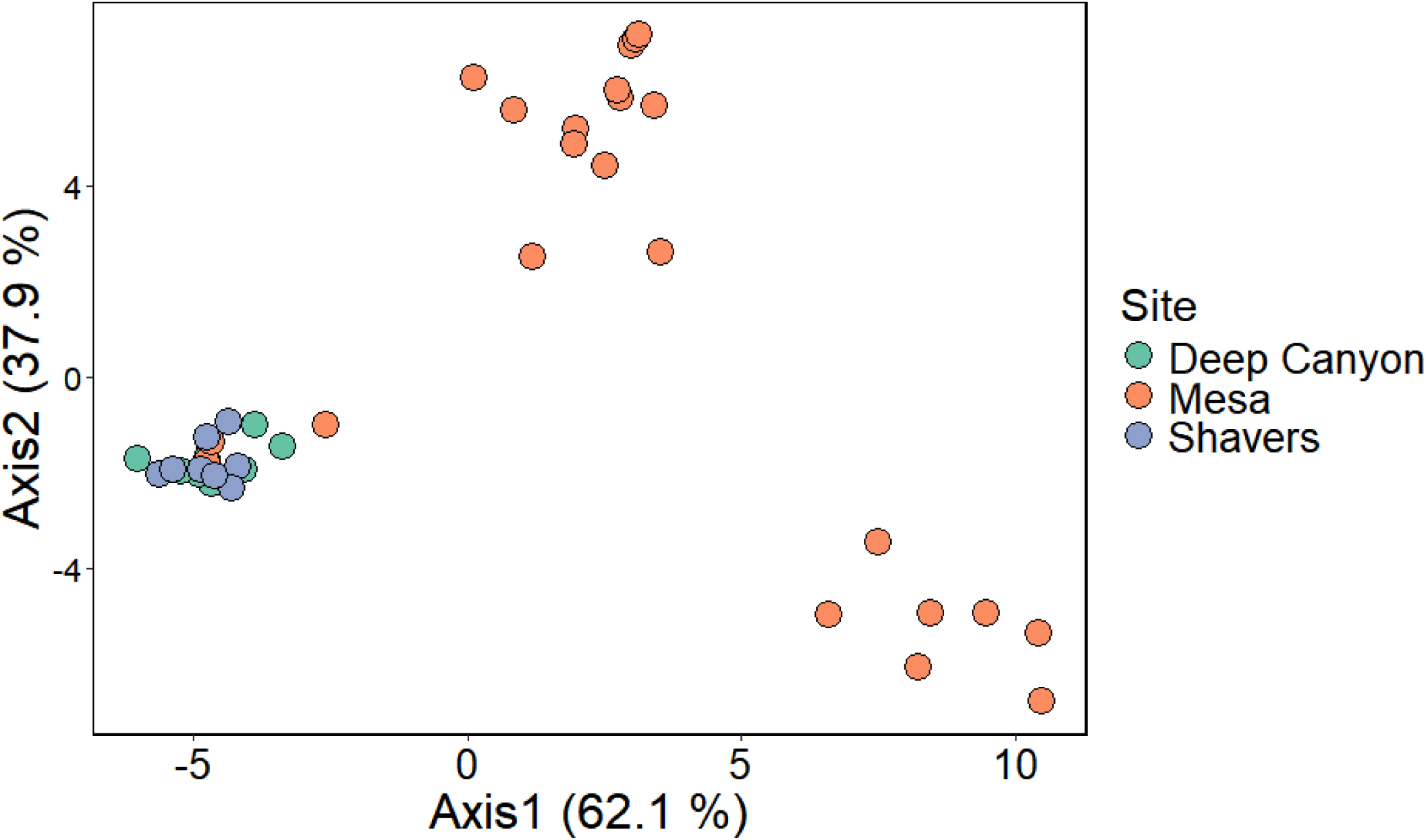
Scatterplot of discriminant functions axes 1 and 2 from the Discriminant Analysis of Principal Component (DAPC) analysis. Each point represents an individual tortoise, and colors correspond to the Deep Canyon, Mesa, and Shavers Valley subpopulations.

#### Estimation of recent migration rates

Parameter estimates obtained from three independent runs of *BA3-SNPs* produced nearly identical results thereby confirming that convergence had been reached. Estimates of recent migration rates (*m_ij_*) from only the first run are shown in Table 1. The recent migration rates into the Mesa subpopulation from the Deep Canyon and Shavers Valley subpopulations were both less than the *a priori* threshold value of *m* = 0.02, which suggests that the Mesa subpopulation is genetically isolated from the other subpopulations (Table 1). Although both estimates of *m* into the Shavers Valley subpopulation from the other two subpopulations were 0.03—and thus exceeded the threshold value, none of the sampled tortoises from the Shavers Valley subpopulation were inferred to be recent migrants from the other two subpopulations (Table 2). In contrast, both recent migration rates into the Deep Canyon subpopulation from the Mesa and Shavers Valley subpopulations (*m* = 0.17 and *m* = 0.08, respectively) also exceeded the critical threshold value (Table 1), but appear to be valid because seven second-generation migrant tortoises were detected in our Deep Canyon subpopulation sample—five tortoises with Mesa ancestry and two with Shavers ancestry (Table 2).

**Table 1.**
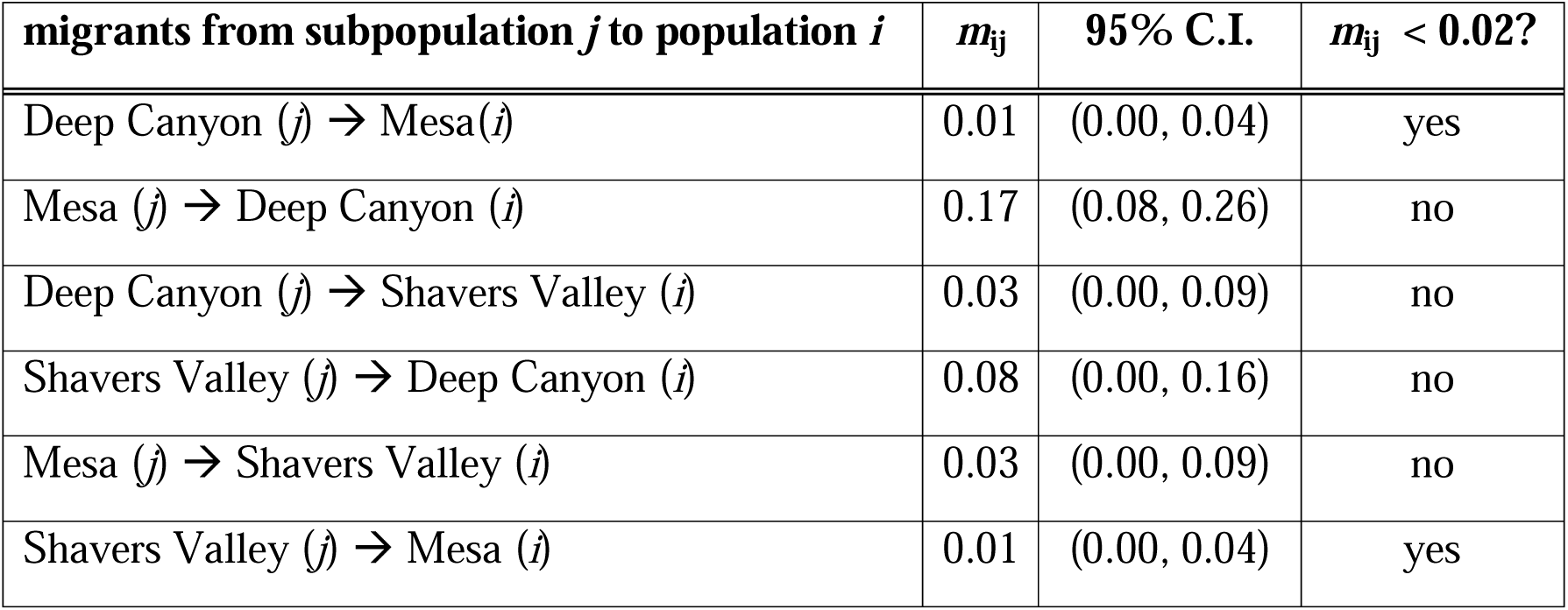
Recent migration rates among the Deep Canyon, Mesa, and Shavers Valley subpopulations. The values for *m*_ij_ are the posterior means for the six migration rate parameters. The parameter *m*_ij_ is defined as the fraction of individuals in subpopulation *i* that are migrants from subpopulation *j*, whereas 95% Credible Interval (C.I.) is the approximate 95% C.I. for each posterior mean. Subpopulation *i* is assumed to be demographically independent of subpopulation *j* when the mean *m*_ij_ < 0.02. Arrows indicate direction of migration.

**Table 2.** Migrant ancestries inferred for the sampled tortoises in this study. Second-generation migrants have one migrant parent and one non-migrant parent. n/a = not applicable.

| **Tortoise I.D.** | **Ancestry** | **Source Subpopulation** | **Probability** |
| --- | --- | --- | --- |
| Deep Canyon 1 | second-generation migrant | Mesa | 1.000 |
| Deep Canyon 7 | second-generation migrant | Mesa | 1.000 |
| Deep Canyon 8 | second-generation migrant | Shavers Valley | 1.000 |
| Deep Canyon 9 | second-generation migrant | Mesa | 1.000 |
| Deep Canyon 10 | second-generation migrant | Mesa | 1.000 |
| Deep Canyon 13 | non-migrant | n/a | 1.000 |
| Deep Canyon 16 | non-migrant | n/a | 1.000 |
| Deep Canyon 17 | second-generation migrant | Mesa | 1.000 |
| Deep Canyon 18 | second-generation migrant | Shavers Valley | 1.000 |
| Mesa 1 | non-migrant | n/a | 1.000 |
| Mesa 2 | non-migrant | n/a | 1.000 |
| Mesa 3 | non-migrant | n/a | 1.000 |
| Mesa 4 | non-migrant | n/a | 1.000 |
| Mesa 6 | non-migrant | n/a | 1.000 |
| Mesa 7 | non-migrant | n/a | 1.000 |
| Mesa 14 | non-migrant | n/a | 1.000 |
| Mesa 21 | non-migrant | n/a | 1.000 |
| Mesa 25 | non-migrant | n/a | 1.000 |
| Mesa 32 | non-migrant | n/a | 1.000 |
| Mesa 33 | non-migrant | n/a | 1.000 |
| Mesa 36 | non-migrant | n/a | 1.000 |
| Mesa 39 | non-migrant | n/a | 1.000 |
| Mesa 43 | non-migrant | n/a | 1.000 |
| Mesa 47 | non-migrant | n/a | 1.000 |
| Mesa 61 | non-migrant | n/a | 1.000 |
| Mesa 69 | non-migrant | n/a | 1.000 |
| Mesa 147 | non-migrant | n/a | 1.000 |
| Mesa 148 | non-migrant | n/a | 1.000 |
| Mesa 150 | non-migrant | n/a | 1.000 |
| Mesa 151 | non-migrant | n/a | 1.000 |
| Mesa 152 | non-migrant | n/a | 1.000 |
| Mesa 161 | non-migrant | n/a | 1.000 |
| Shavers Valley 1 | non-migrant | n/a | 1.000 |
| Shavers Valley 2 | non-migrant | n/a | 1.000 |
| Shavers Valley 7 | non-migrant | n/a | 1.000 |
| Shavers Valley 8 | non-migrant | n/a | 1.000 |
| Shavers Valley 12 | non-migrant | n/a | 1.000 |
| Shavers Valley 13 | non-migrant | n/a | 1.000 |
| Shavers Valley 28 | non-migrant | n/a | 1.000 |
| Shavers Valley 31 | non-migrant | n/a | 1.000 |

#### Estimation of contemporary effective population sSizes

The program *CurrentNe* estimated *N*_eLD_ for the Deep Canyon subpopulation to be only five breeding adult tortoises with the 90% confidence interval (C.I.) ranging from 5-8 individuals. For the Mesa subpopulation, *N*_eLD_ was estimated to be 13 adults with a C.I. ranging from 11-17 individuals while the Shavers Valley subpopulation *N*_eLD_ was estimated to be 36 individuals (90% C.I. 16-80).

## Discussion

### Genetic structure of desert tortoises in the Coachella Valley region

The Mesa, Deep Canyon, and Shavers Valley subpopulations of desert tortoises exhibited weak population structure according to our model-based *fastStructure* analyses. Although both the *fastStructure* and DAPC analyses suggested that *K* = 3 best explains the data, the three defined clusters did not equate to the Mesa, Deep Canyon, and Shavers Valley subpopulations; instead, they corresponded to a single cluster that comprised the Deep Canyon and Shavers tortoises plus two genetically divergent Mesa clusters. Thus, these results suggested that the Deep Canyon and Shavers Valley subpopulations were more closely related to each other than either was to the Mesa subpopulation, and that there was little structure within the Deep Canyon/Shavers Valley cluster. In contrast, we observed substantial structure within the Mesa subpopulation that may be the result of historical gene flow from another tortoise subpopulation. Support for this interpretation of our results comes from the study by Lovich et al. (2020), who demonstrated that the Mesa tortoises were admixed with tortoises from the Western Mojave (WM) recovery unit (USFWS 1994).

Despite the evidence for gene flow from the WM recovery unit into the Mesa subpopulation, environmental barriers may have limited genetic interchanges between these tortoise subpopulations over evolutionary time. One of us (Lovich) observed a few tortoises in the Morongo Valley, which is ∼15 airline miles north of the Mesa tortoises. But it is not clear if those individuals lived there or were migrants or released captives from the adjacent Yucca Valley and Joshua Tree areas. The presence of a steep elevational/environmental gradient between the Mesa site and Morongo Valley, which is a transitional zone between low elevation Colorado Desert and higher elevation Mojave Desert, may have limited the amount of genetic mixing between subpopulations owing to the differentiating effects of divergent natural selection (see Kawecki and Ebert 2004; Samuk et al. 2017; Harringmeyer and Hoekstra 2022). Also, the Little San Bernardino Mountains presumably acted as a barrier to gene flow between Mesa and any tortoise subpopulations that occurred in Morongo Valley before modern development altered their habitat. Loss of tortoise habitat in the Morongo Valley in recent decades due to urbanization has probably reduced further or even cut-off gene flow between Mesa and WM tortoises.

#### Recent migration rates among the three tortoise subpopulations

Given the apparent isolation of the Deep Canyon, Mesa, and Shavers Valley subpopulations, we were surprised to see that only two of the six migration rate estimates indicated negligible amounts of migration over the past couple tortoise generations. Recent migration rates into the Mesa subpopulation from the Deep Canyon and Shavers Valley subpopulations were 0.01 and therefore less than the *a priori* threshold value (i.e., *m* < 0.02) indicating that the Mesa subpopulation is demographically independent of the other two sampled subpopulations. In contrast, estimates of *m* into the Deep Canyon subpopulation from the Shavers Valley and Mesa subpopulations were 0.08 and 0.17, respectively, and thus both were well above the threshold, while recent gene flow into the Shavers Valley subpopulation from Deep Canyon and Mesa subpopulations were 0.03 each and therefore slightly exceeded the threshold.

The *BayesAss/BA3-SNPs* program estimates recent migration rates by identifying first-and second-generation migrants in a sample of individuals (Wilson and Rannala 2003). Accordingly, the basis of our significant gene flow estimates into the Deep Canyon subpopulation becomes evident once we see that the program identified seven second-generation tortoises (of nine sampled) at Deep Canyon: two were inferred to have one parent from the Shavers Valley subpopulation while two others had one parent that originated in the Mesa subpopulation. However, all eight of the tortoises we sampled in the Shavers Valley subpopulation were identified as non-migrants, which suggests that both significant estimates of *m* (i.e., 0.03) going into the Shavers Valley subpopulation could be spurious artifacts, perhaps caused by a lack of signal due to inadequate sample size and/or interactions with the prior. Forester et al. (2022a) stated without justification that sample sizes of at least 30 individuals per population or subpopulation are needed to detect migrants using genetic approaches such as *BayesAss3/BA3-SNPs*. Contrary to this assertion, our results suggest that useful results can be obtained from this program even with low sample sizes. Indeed, although we only sampled eight and nine tortoises from Shavers Valley and Deep Canyon, respectively, we nonetheless detected multiple individuals having migrant ancestries in our sample. Given how *BayesAss3/BA3-SNPs* estimates recent migration rates, our success in detecting migrants at Deep Canyon with such a small number of tortoises could be attributable to the fact that we sampled ∼70% of the subpopulation (see below). Thus, what matters more may not simply be the total number of individuals sampled, but the proportion of a subpopulation or population that is sampled. Further work is needed to determine whether the procedures and sample sizes we used can reliably identify MUs using such a low migration rate threshold of 0.02. Although our results suggest that only the Mesa and Shavers Valley subpopulations represent MUs, below we argue that the Mesa and Deep Canyon subpopulations merit designations as MUs while the MU-status of the Shavers Valley subpopulation is uncertain for now.

#### Is the Deep Canyon subpopulation part of a metapopulation or is it an MU?

How did desert tortoises migrate to Deep Canyon from the Shavers and Mesa subpopulations? First, we will consider the migrant(s) from the Shavers Valley subpopulation, which is located ∼50 airline km away from Deep Canyon (Figure 1). Prior to agricultural development in the Coachella Valley, which began about 1900 (Meyer and van Schilfgaard 1984), the central part of the valley was covered by vast wind-blown sand dune fields (Beatley 1992). Such dune fields would have likely prevented desert tortoises from crossing the Coachella Valley since it is poor tortoise habitat. Since then, the valley has been developed into agricultural fields and urban areas, and a railroad track was built down through the valley towards the Salton Sea. Railroads are known to act as barriers to tortoise and turtle movements (Pelletier et al. 2005; Kornilev et al. 2006; Iosif 2021; Platt et al. 2018). Thus, the only plausible explanation is that humans transplanted the tortoises from Shavers Valley to Deep Canyon.

There are numerous scenarios by which human-mediated transplanting of tortoises to Deep Canyon could have occurred. For example, military personnel who were training near the Shavers Valley study site just before the start of World War II (Henley 2000) could have moved tortoises to Deep Canyon as there was a military training camp based at the mouth of Deep Canyon. It is also possible that native Americans living in the desert (Schneider and Everson 1989) could have moved some tortoises around to different desert sites though the nature of their interactions with desert tortoises is not known to us. It is also possible that around the time when the Living Desert Reserve was established next to Deep Canyon in 1970 that one or more tortoise(s) originally obtained from the Shavers Valley area were released into Deep Canyon.

Still another scenario could be that captive tortoises from Shavers Valley were released in or near the Deep Canyon population. For example, between 1971 and 1972, 65 tortoises were released south of Deep Canyon in Anza Borrego Desert State Park (ABDSP), and some of them reproduced (Luckenbach 1982). A population persists at ABDSP (Manning 2018; Manning et al. 2022) although Luckenbach considered those tortoises to be outside their natural range. The distance between Deep Canyon and ABDSP is over 40 km, and the terrain includes high mountains, so it is unlikely that tortoises moved between the sites. It is important to emphasize that all these scenarios are speculative, as we have no specific evidence for any of them at this time.

The various human-mediated translocation hypotheses mentioned above can also explain how tortoise(s) from the Mesa site arrived at Deep Canyon and thus we will now consider the alternative hypothesis—that unassisted or natural migration occurred instead. As pointed out before, the straight-line distance between the Mesa and Deep Canyon sites is ∼30 km (Figure 1). However, the actual distance a tortoise would need to travel between sites is closer to 40-50 km according to Google Earth. Hypothetically, there are only two routes that a tortoise could have taken to reach Deep Canyon from the Mesa site. In one possible (“lower elevation”) route, tortoise(s) could have followed the lower desert slopes and alluvial fans of the San Jacinto and Santa Rosa Mountains until reaching Deep Canyon. This route would only have been a possible option prior to the extensive urbanization of the Palm Springs area, which began by at least the 1940s (USGS Toro Peak, 1941), because the natural habitat along this route has largely been replaced by human infrastructure (roads, buildings, etc.). In another possible (“higher elevation”) route, tortoise(s) might have reached Deep Canyon by climbing up and down the numerous steep canyons above the lower elevation route. Owing to the ruggedness of the terrain along this route with the possibility that at least one of the canyons might have been impassable to tortoises, we think it is unlikely that there could have been much if any migration of tortoises along this route in historical times. Given the improbability of tortoises migrating from the Shavers and Mesa subpopulations to Deep Canyon along with the unlikelihood there will be no unsanctioned human-assisted translocations to Deep Canyon in the future, we regard this subpopulation as an MU.

#### Is the Shavers Valley subpopulation part of a metapopulation or is it an MU?

Although our migration rate analyses suggest that the Shavers Valley subpopulation may also be defined as an MU, it is premature to proceed with this designation until further studies of the EC recovery unit are conducted. This is because the “Shavers Valley subpopulation,” as we have defined it, likely represents a fraction of a much larger population that occupies the eastern Colorado Desert (i.e., EC recovery unit). Tortoise habitat extends far beyond the Shavers Valley site—into Joshua Tree National Park to the north, the Orocopia and Chocolate Mountains to the south, and the Chuckwalla Mountains to the east. Admixture plots in Sánchez-Ramirez et al. (2018) and Lovich et al. (2020) showed that tortoises throughout the eastern Colorado Desert in California (i.e., east of the Coachella Valley) formed a single genetic cluster, which may reflect recent or on-going genetic connectivity among these areas. New surveys should be conducted throughout the range of the EC recovery unit to collect additional genetic samples so that all MUs within this recovery unit can be defined.

#### Evaluating the local *N*_eLD_ estimate for each desert tortoise MU

The *N*_eLD_ estimates for the Deep Canyon, Mesa, and Shavers Valley subpopulations were 5, 13, and 36 adults, respectively. If these estimates are correct, then our findings suggest that the Deep Canyon and Mesa MUs have high inbreeding rates (*N*_eI_ < 50) and are in danger of losing excessive amounts of additive genetic variation over the long term (*N*_eAV_ < 500). Due to the uncertainty surrounding the MU status of the Shavers Valley subpopulation, it not yet possible to determine the genetic health of this subpopulation though as we discuss below it likely has a much larger *N*_e_ than the other two subpopulations.

How trustworthy are our *N*_eLD_ estimates given that they are used as proxies for *N*_eI_ and *N*_eAV_? Although little is known about the effects of sample size on the estimation of *N*_eLD_, we will first consider the sample sizes we used to estimate *N*_eLD_ for each subpopulation. The *N*_eLD_ estimate for the Mesa MU is based on a sample of 23 individuals, most of which were adults.

Considering that the census size for this subpopulation was around 32 adults around the time when we obtained our genetic samples (Lovich et al. 2011), it seems reasonable to think that we obtained a *N*_eLD_ estimate that is close to the true *N*_e_ value, especially since this population is likely closed. Likewise, the *N*_eLD_ estimate for the Deep Canyon MU is based on nine of the estimated total census size (*N*) of 13 adults (70%) for this subpopulation. However, our estimate of *N*_eLD_ for the Deep Canyon MU may be confounded by the recent influx of migrant tortoises into the Deep Canyon subpopulation, an issue we will discuss below.

As alluded to above, there is good reason to think that our *N*_eLD_ estimate for the Shavers Valley subpopulation is a substantial underestimate of the true *N*_e_. All eight of the samples we obtained from this subpopulation were obtained from a small plot in the foothills of the Cottonwood Mountains on the southern border of Joshua Tree National Park. However, as we pointed out above it is highly probable that the Shavers Valley subpopulation is a subset of a larger tortoise subpopulation that exists in surrounding areas (e.g., Lovich et al. 2014). Another sign that we did not sample tortoises from a large enough area around the Shavers Valley site is that the *N* estimate for the Shavers Valley subpopulation, which is 30 individuals (Lovich and Cummings unpublished data), is smaller than our *N*_eLD_ estimate of 36 adults. If the Shavers Valley subpopulation was truly closed, and we had adequately sampled from it, then we would expect the *N*_e_/*N* ratio to be much less than one. This is because the *N*_e_/*N* ratio for a polygamous species like the desert tortoise should be less than one (see Waples 2024). Yet the *N*_e_/*N* ratio for the Shavers Valley subpopulation is 1.2, which indicates something may be amiss with our estimate(s)—most likely both numbers are underestimates, and *N*_e_ should be much smaller than *N*. Thus, additional sampling should be done throughout the eastern Colorado Desert in California so that we can better understand the actual sizes of *N*_e_ and *N* for the Shavers Valley subpopulation.

Although we consider the Deep Canyon subpopulation to be currently closed to immigration, evidence of recently transplanted individuals from the Mesa and Shavers Valley subpopulations into this subpopulation might raise doubts about the validity of our local *N*_eLD_ estimate for the Deep Canyon MU given that even slight levels of migration can result in *N*_eLD_ values that are underestimates of *N*_eI_ and *N*_eAV_ (Ryman et al. 2019). However, because these “migration” events appear to have been one-time or otherwise exceedingly rare occurrences rather than constant migration events every generation (on average) over evolutionary time, we suspect that our *N*_eLD_ estimate for this subpopulation probably closely reflects the correct value for the true local *N*_e_. We can attempt to corroborate our genetics-based estimate of *N*_e_ using an alternative approach to estimating local *N*_e_.

If an estimate for *N* is available along with a reasonable empirical estimate for *N*_e_/*N* ratio, then one can extrapolate *N*_e_ (Palstra and Ruzzante 2008; Luikart et al. 2010; Palstra and Fraser 2012; Waples 2024). Given our estimates of *N* for the Deep Canyon (13 adults) and Mesa (32 adults) MUs (Lovich, Cummings, and Tracy unpublished data), we can couple the *N* estimate for the Mesa MU with its estimated *N*_e_ of 13 adults to obtain an *N*_e_/*N* ratio of 0.406, a reasonable estimate given that this species has a polygamous mating system. Then using the Mesa *N*_e_/*N* ratio with an *N* of 13 for the Deep Canyon MU gives us an *N*_e_ = 5 adults for the Deep Canyon MU, which is the same as our *N*_eLD_ estimate. Note that even if we instead use the median *N*_e_/*N* value of 0.231 from the multitude of studies that were vetted by Palstra and Fraser (2012), the estimated *N*_e_ for the Deep Canyon MU would be three adult tortoises, which is negligibly different from our other estimate in the context of the 50/500 rule.

#### Should the Mesa and Deep Canyon MUs be genetically rescued?

The genetic rescue hypothesis holds that the genetic diversity of small, inbred populations can be beneficially augmented through human-mediated transplants of individuals from other populations (Ingvarsson 2001). This idea grew out of observations from experimental studies whereby investigators discovered that the introduction of just one to few individuals to small, inbred populations could bring fitness benefits to those populations (Spielman and Frankham 1992; Madsen et al. 1999; Ball et al. 2000; Newman and Tallmon 2001). However, genetic rescue is not without its risks, as some workers have voiced concerns that this practice might cause more harm than good to populations on the brink. One concern is that if the introduced individual(s) are from a population that is too evolutionarily divergent from the population to be rescued, then the result could end up being a reduction in population fitness, a phenomenon labeled “outbreeding depression” (Goldberg et al. 2005; Tallmon et al. 2004; Murphy et al. 2007; Edwards and Berry 2013; MacDonald et al. 2025). The authors in these studies advocated a more cautious approach by first conducting small-scale controlled experiments to determine the likelihood for success if a genetic rescue program were to be initiated. Such an approach, however, is not feasible for desert tortoises because they have generation times that are about two decades in length and thus it would take many decades to obtain the information needed, which may come too late to make a positive difference.

The data in this study show that the Mesa, Deep Canyon, and Shavers Valley subpopulations of desert tortoises are not evolutionarily divergent from each other. This is not surprising given that all three subpopulations live in habitats that are similar climatically and thus they all may have similar adaptations for coping with local environmental conditions. This may help explain how the migrant tortoises at Deep Canyon were able to successfully breed with non-migrant tortoises. It is too early to deem previous introductions as being successful in a genetic rescue sense, but the population should continue to be monitored to determine if the tortoises of mixed ancestry are able to produce viable offspring. Nonetheless, the results of this study suggest that the Deep Canyon and Mesa subpopulations might be good candidates for genetic rescue.

## Acknowledgements

WBJ is grateful for assistance provided by Wei Zhang, Holly Clark, and Clay Clark of the UCR Genomics Core, Emerson Jacobson of the UCR HPCC, Paige Meija and Joel Sachs of the EEOB Department at UCR, and Jason Boone at Floragenex, Inc. Al Muth and Chris Tracy provided accommodations at the Philip L. Boyd Deep Canyon Research Center of the University of California, Riverside (doi:10.21973/N3V66D), during our research. Wildlands Inc. provided access to their land near the Orocopia Mountains for tortoise surveys and Michael Vamstad of the National Park Service did the same for surveys in southern Joshua Tree National Park. Research was conducted under permits from the U.S. Fish and Wildlife Service, the Bureau of Land Management, the National Park Service, and the California Department of Fish and Wildlife. We are grateful to the Institutional Animal Care and Use Committee of Northern Arizona University for reviewing and approving our research procedures.

## Data availability

We deposited the original Illumina sequence reads and SNP genotype datafiles used in analyses in the Dryad repository under DOI: xxxxxxx.

